# Rapid *Mycobacterium tuberculosis* spoligotyping from uncorrected long reads using Galru

**DOI:** 10.1101/2020.05.31.126490

**Authors:** Andrew J. Page, Nabil-Fareed Alikhan, Michael Strinden, Thanh Le Viet, Timofey Skvortsov

## Abstract

Spoligotyping of *Mycobacterium tuberculosis* provides a subspecies classification of this major human pathogen. Spoligotypes can be predicted from short read genome sequencing data; however, no methods exist for long read sequence data such as from Nanopore or PacBio. We present a novel software package Galru, which can rapidly detect the spoligotype of a *Mycobacterium tuberculosis* sample from as little as a single uncorrected long read. It allows for near real-time spoligotyping from long read data as it is being sequenced, giving rapid sample typing. We compare it to the existing state of the art software and find it performs identically to the results obtained from short read sequencing data. Galru is freely available from https://github.com/quadram-institute-bioscience/galru under the GPLv3 open source licence.

## Introduction

Tuberculosis (TB), caused mainly by Mycobacterium tuberculosis, is a disease of global importance involved in 1.5 million fatalities in 2018 (WHO | Global tuberculosis report 2019). There is an increasing prevalence of drug-resistant TB.

Over the past 30 years, the global effort in surveillance and treatment has been supported by a number of molecular genotyping tools. These methods use a variety of targets, such as the length of tandem repeat patterns (MIRU-VNTR), or single nucleotide variants from whole genome sequencing (Meehan et al., 2019). One method in particular, “spoligotyping” (a portmanteau of spacer oligonucleotide typing) detects the variation within the clustered regularly interspersed short palindromic repeats (CRISPR) locus (Kamerbeek et al., 1997). This locus, sometimes referred to as the Direct Repeat locus in the context of M. tuberculosis, contains an alternating series of a 36 base-pair repeating sequence and a 35 to 41 base-pair variable “spacer” sequence (Hermans et al., 1991). This pattern, particularly the presence or absence of 43 specific spacer sequences, varies across the species and can be used to infer the underlying transmission or evolutionary history of M. tuberculosis. The hybridization pattern, the “spoligotype”, can be summarised as a 43-digit binary code denoting the presence (1) or absence (0) of a particular spacer. The spoligotype can be further shortened into a hexadecimal or octal code (Dale et al., 2001).

High throughput sequencing is being increasingly accepted as the preferred method of identifying microbial pathogens, particularly for national surveillance and outbreak investigation (Nadon *et al.*, 2017; Pérez-Losada *et al.*, 2018). These diagnostics are now approaching turnaround times of hours by direct sequencing of metagenomic samples, or operate at a competitive cost often undercutting traditional microbiology methods by bulk sequencing samples (Gardy and Loman, 2018). This is a marked improvement over even relatively recent advances such as RFLP typing which can still take approximately 26 days for a result (Gori *et al.*, 2005). Taxonomic identification to the species (or finer level) is a critical diagnostic step, which is usually calculated by detecting a small set of curated genomic markers with programs such as MIDAS (Nayfach *et al.*, 2016) or MetaPhlan2 (Truong *et al.*, 2015); or by aligning reads to a set of reference genomes, the approach implemented in Kraken (Wood and Salzberg, 2014) and MEGAN (Huson *et al.*, 2016). Either case requires surveying the entire genome. This may not always be required in *M. tuberculosis,* as spoligotyping focuses on a singular locus rich with genetic diversity (the CRISPR locus) and has already proven effective.

CRISPR loci can be resolved within a single long read such as produced by Sanger sequencing, Illumina TruSeq SLR (Nasko *et al.*, 2019; Lam and Ye, 2019), or from PacBio or Nanopore platforms. Use of long-read sequencing on metagenomic samples offers several notable advantages such as omitting PCR steps, which can introduce contamination and complexity as well as limiting the range of information available. Yet, there was no tool available that utilises the latest advancements in long read sequencing to produce familiar spoligotyping results. Here we provide a new twist on the existing typing protocols with *Galru*, a Python 3 program that defines spoligotyping of *M. tuberculosis* directly from uncorrected long reads, and is available under the open source licence GNU GPL 3 from https://github.com/quadram-institute-bioscience/galru.

## Methods

### Implementation

The basic method is to identify reads containing complete CRISPR-associated systems for *M. tuberculosis*, which get mapped to a database of known spacers for spoligotyping. This enables identification of the spoligotype from a single read.

### Database of CRISPR regions

Annotated complete reference genomes were downloaded from NCBI (RefSeq, accessed 2020-05-24) for *M. tuberculosis*. CRISPR regions, spacers and repeats, were identified within the genomes using minced (Skennerton, 2019) (v0.4.0), and the full nucleotide regions were extracted using bedtools (Quinlan and Hall, 2010) (v2.27.1). The nucleotide sequences were assigned a unique identifier and added to a FASTA file. These sequences were subsequently clustered using CD-HIT (Li and Godzik, 2006; Fu *et al.*, 2012) (version 4.8.1) at 99% nucleotide identity to remove near identical sequences. This has the effect of reducing the size of the database and speeds up the identification of candidate reads. A prebuilt database is provided with Galru along with scripts to build new databases for other species.

### Spoligotyping method

The input to Galru is a FASTQ or FASTA file of uncorrected long reads. These reads can be streamed in as they are generated, such as allowed for with the Nanopore sequencing technology and can optionally be compressed. Assemblies can also be used as input. The reads are first mapped to the CRISPR regions database using minimap2 v2.17 (Li, 2018), with the parameters set by the sequencing technology specified by the user. Any reads which map to any CRISPR region, regardless of mapping score, are passed to the next stage, with all other reads filtered out. This reduces false positives and speeds up the computation. A modified reciprocal blast (v2.9.0) (Camacho *et al.*, 2009) is performed between the spoligotyping spacers and the filtered reads containing the CRISPR region. By default a single mismatch is allowed within the 25 base spacers used for typing, which can be modified at runtime to allow for a stricter or looser assignment. If a spacer sequence is found to be present in a single read, it is said to be present within the genome. The spoligotype is outputted in a binary format, with 0 for absent and 1 for present, as is community practice.

## Samples

The largest public dataset of Nanopore data for *M. tuberculosis* contains 5 samples which were sequenced using short and long read sequencing technologies (Hunt *et al.*, 2019). The full list of accession numbers is available in Table 1. All of the data was downloaded from the European Nucleotide Archive.

**Table 1:** Accession numbers for sample datasets used for validation.

| Sample Accession | Illumina Accession | ONT Accession |
| --- | --- | --- |
| ERS3036286 | ERR3077901 | ERR3078032 |
| ERS3036287 | ERR3077902 | ERR3078033 |
| ERS3036288 | ERR3077903 | ERR3078034 |
| ERS3036289 | ERR3077904 | ERR3078035 |
| ERS3036290 | ERR3077905 | ERR3078036 |

## Results

To validate the accuracy of Galru, the short read Illumina data was provided to SpoTyping (v1.5) (Xia *et al.*, 2016), the state of the art software application for spoligotyping from short read data. SpoTyping was provided with short read Illumina data and Galru was provided with long read Nanopore data as shown in Table 2. In every case the spoligotypes identified were identical. The short read Illumina data has a higher per base quality and a higher coverage when compared to the long read Nanopore data. The running time for both SpoTyping and Galru was under 10 seconds for each sample.

**Table 2:**
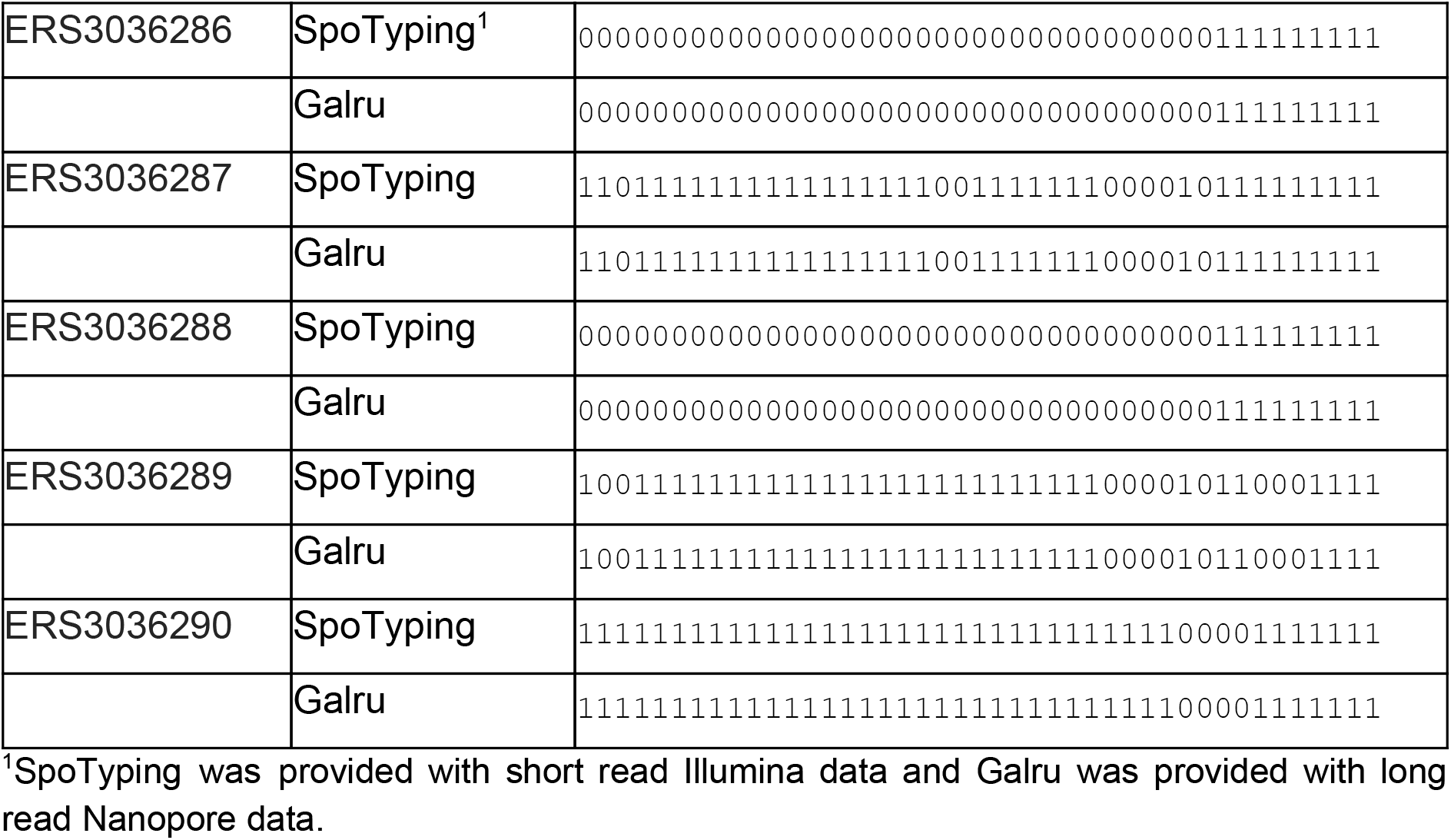
The spoligotyping results for each sample using SpoTyping and Galru.

## Discussion

As DNA sequencing technologies mature, whole genome sequencing becomes a reliable and versatile alternative for characterisation of bacterial isolates, being gradually accepted and adopted for clinical diagnostics, epidemiological surveillance and outbreak investigation. Whole genome analysis provides the maximum information about the isolate, but requires genome assembly and/or mapping of reads to a reference genome, which makes it computationally intensive and time-consuming; therefore, direct analysis of specific loci represents a more attractive alternative, without substantial resolution losses.

Among the loci that can be used for such an analysis, CRISPR have been quite popular for typing a number of other pathogenic bacteria, including *Yersinia pestis and Salmonella* (Tang *et al.*, 2019). This is exemplified in the *M. tuberculosis* complex with the implementation and continued use of spoligotyping. Nevertheless, due to specific structural organisation of CRISPR arrays, their accurate assembly might present a problem. Third generation sequencing systems (Nanopore/PacBio) produce reads long enough to completely cover the whole CRISPR region in a single read, thus avoiding the genome assembly step and circumventing problems arising from its use. However, the per base quality of long read sequencing technologies is lower than for short read sequencing, so different solutions are required to cater for this error model.

We have implemented a software that can reliably predict spoligotypes in long read sequencing data for *M. tuberculosis.* It is fast and just as accurate as spoligotyping from short read sequencing technologies as we showed by comparing it to the state of the art software SpoTyping. SpoTyping however cannot handle uncorrected long read sequencing data, it must first be assembled *de novo*, and polished, a resource intensive task from long read sequencing data, requiring substantial coverage. Galru does not require *de novo* genome assembly and in fact can perform typing from a single long read. Nanopore sequencers can produce basecalled data in near real-time, thus allowing for reads to be analysed by Galru as they are produced, providing results rapidly from a minimal amount of sequencing information. It is possible to use Galru to analyse metagenomic datasets given its minimal coverage requirements, opening the door for spoligotyping directly from uncultured clinical samples, further reducing the time from swab to result.

## Conclusion

Spoligotyping remains a valuable tool for the continued surveillance of *M. tuberculosis*. Galru provides a stepwise improvement by allowing rapid spoligotyping directly from long read sequencing. It is fast and accurate, requires a minimal amount of information to produce a spoligotype, and allows for near real-time typing when used to process sequencing data as it is produced by a Nanopore sequencer. Galru was written in Python and is available under the open source licence GNU GPL 3 from https://github.com/quadram-institute-bioscience/galru. It includes unit tests for validation of the software and is easily installable using pip, conda or Galaxy.

## Acknowledgements

Authors wish to thank Zamin Iqbal for identifying the *M. tuberculosis* dataset.

## Author contributions

All authors have read this manuscript and consented to its publication. AJP wrote the software and developed the method. NFA contributed to the software and method. TS, NFA, MS performed the literature review. AJP, NFA, MS and TS wrote the manuscript. TLV packaged the software for ease of installation. AJP and TS provided project management and oversight. AJP secured funding for the project.

